# Oxazolidinone antibiotics exhibit potent in vitro activity against Gram-positive bacterial endophthalmitis isolates

**DOI:** 10.64898/2026.10.05.756736

**Authors:** Marwa G. Rahim, Emily A. Young, Eric G. Romanowski, Andrew W. Eller, Robert M. Q. Shanks

## Abstract

**Purpose:** Vancomycin-resistant Gram-positive organisms represent a potential therapeutic challenge in endophthalmitis. To evaluate alternative treatment options, the FDA-approved oxazolidinones linezolid (LZD) and tedizolid (TZD) were tested against a collection of Gram-positive bacterial endophthalmitis isolates.

**Methods:** Minimum inhibitory concentrations (MICs) were determined using epsilometers (Etest strips) for coagulase negative staphylococci (CoNS, n=20), *Staphylococcus aureus* (n=18), *Bacillus* species (n=10), *Enterococcus* species (n=5), and *Streptococcus pneumoniae* (n=1). Available CLSI and EUCAST breakpoints were applied to determine susceptibility.

**Results:** TZD exhibited a >5-fold reduction in median MIC compared with LZD across all organisms tested (p<0.0001). MIC_90_ values for CoNS were 2.0 µg/ml for LZD and 0.25 µg/ml for TZD. For *S. aureus*, MIC_90_ values were 1.5 µg/ml (LZD) and 0.19 µg/ml (TZD). For *Bacillus* spp., MIC_90_ values were 1.0 µg/ml (LZD) and 0.25 µg/ml (TZD). Among *Enterococcus* species, the MIC range was 0.5-4.0 µg/ml (LZD) and 0.064-0.75 µg/ml (TDZ). The single *S. pneumoniae* strain had a MIC of 0.5 for LZD and 0.064 for TZD. All isolates were susceptible to TZD. One *Enterococcus* isolate was non-susceptible to LZD; the remaining isolates were susceptible.

**Conclusions:** LZD and TZD demonstrated in vitro activity against Gram-positive bacterial endophthalmitis isolates, with TZD displaying consistently lower MICs than LZD. These findings suggest that oxazolidinones may represent potential alternatives when vancomycin cannot be used, including infections caused by vancomycin-resistant Gram-positive organisms. Given their excellent oral bioavailability, favorable safety profile, and reported anti-virulence effects in certain pathogens, further evaluation of oxazolidinones for the treatment of intraocular infections is warranted.

## Introduction

Bacterial endophthalmitis is among the most vision-threatening infectious diseases of the eye and remains an ophthalmic emergency because irreversible vision loss can occur within hours of symptom onset. In the United States and many other countries this is frequently caused by Gram-positive bacteria such as coagulase negative staphylococci, though it can be caused by a variety of organisms ^1,2,3^. Endophthalmitis most commonly develops following intraocular surgery but may also occur due to ocular trauma, intravitreal injection, or hematogenous spread from systemic infections. Some pathogens, including *Bacillus cereus* and toxin-producing *Enterococcus* species, can cause profound retinal damage and blindness in hours, highlighting the need for rapid antimicrobial intervention ^4,5^.

Because microbiological identification and susceptibility testing require time, treatment of endophthalmitis is typically empiric. Current standard therapy consists of intravitreal antibiotics combinations active against both Gram-positive and Gram-negative bacteria, most commonly vancomycin with ceftazidime ^1,6,7^. This regimen has remained highly effective against most ocular isolates; however, concerns regarding antimicrobial resistance and the limited number of alternative agents available for resistant infections remain. While vancomycin-resistant *Enterococcus* (VRE) are not a common endophthalmitis pathogen, cases have been reported and represent a significant therapeutic challenge ^8,9^. More broadly, recent outbreaks of highly resistant ocular pathogens have highlighted the vulnerability of current treatment paradigms. In 2023, an outbreak of extensively drug-resistant *Pseudomonas aeruginosa* keratitis associated with contaminated artificial tears demonstrated how antimicrobial resistance can render standard empiric therapies ineffective and result in devastating visual outcomes ^10,11^. Such events underscore the importance of evaluating alternative antimicrobial strategies before resistant organisms become established causes of intraocular infection.

Although vancomycin remains a highly effective bactericidal agent against most Gram-positive ocular pathogens, concerns regarding toxicity and stewardship persist ^12,13^. Intercameral vancomycin prophylaxis has been associated with rare but severe retinal complications, including hemorrhagic occlusive retinal vasculitis ^14^. Furthermore, vancomycin is considered a critical therapeutic agent for serious systemic infections caused by methicillin-resistant *Staphylococcus aureus* (MRSA) and other resistant Gram-positive organisms, and antimicrobial stewardship initiatives generally discourage unnecessary use when alternatives are available ^12^.

Oxazolidinones are a class of synthetic antimicrobial agents with potent activity against Gram-positive bacteria ^15^. The two U.S. Food and Drug Administration (FDA)-approved agents, linezolid, and the second-generation compound tedizolid, inhibit protein synthesis by preventing formation of the functional 70S ribosomal initiation complex. Because this mechanism differs from those of other protein synthesis inhibitors, cross-resistance with other ribosome-targeting antimicrobial classes is uncommon. Both agents demonstrate activity against clinically important Gram-positive pathogens, including MRSA and VRE ^16,17^. In addition to antibacterial activity, linezolid has been reported to suppress toxin production and other virulence-associated phenotypes in several Gram-positive bacterial genera ^16,17^. From a therapeutic perspective, linezolid is notable for its excellent oral bioavailability and ability to penetrate tissues including the central nervous system ^18^. This pharmacologic property suggests that it could be used orally or injected.

Despite their FDA approved systemic use, oxazolidinones have not been systematically evaluated against a collection of bacterial endophthalmitis isolates. Before studies of ocular safety, pharmacokinetics, local delivery, or clinical efficacy can be justified, activity against endophthalmitis pathogens must first be established. Therefore, the purpose of this study was to evaluate the in vitro activity of linezolid and tedizolid against a collection of Gram-positive bacterial endophthalmitis isolates and to assess their potential utility as an alternative or adjunctive therapeutic for the treatment of intraocular infection.

## Methods

### Bacterial Strains and Susceptibility Testing Method

De-identified, laboratory confirmed clinical bacterial strains isolated from patients with endophthalmitis presenting to the Department of Ophthalmology at the University of Pittsburgh School of Medicine, in Pittsburgh, PA, USA, were used for *in vitro* susceptibility testing. Isolates of coagulase negative staphylococci (CoNS, n=20), *Staphylococcus aureus* (n=18), *Bacillus* species (n=10), *Enterococcus* species (n=5), and *Streptococcus pneumoniae* (n=1) were isolates stored at -80^°^C and were streaked onto blood agar (tryptic soy agar with 5% sheep erythrocytes, BBL) and incubated overnight at 37^°^C.

The following day, a loopful of bacterial growth was transferred from the blood agar into a glass tube containing 5 mL of phosphate-buffered saline (PBS) and mixed to prepare the bacterial suspension. A sterile dacron-coated swab was immersed in the bacterial suspension, and the swab was used to inoculate Mueller-Hinton agar (BBL) plates (or Mueller-Hinton agar supplemented with 5% sheep erythrocytes for the *Streptococcus* isolate, BBL). The agar surface was streaked in three directions to ensure uniform distribution of the bacteria.

Linezolid and tedizolid epsilometer strips (Liofilchem) were then placed on opposite sides of the inoculated agar using tweezers. The strips were inverted so the highest concentrations of antibiotics were opposite from each other. Flame sterilization was performed throughout the procedure to maintain aseptic technique. The plates were incubated at 37°C for approximately 24 hours. Following incubation, the minimum inhibitory concentration (MIC) values for linezolid and tedizolid were read and recorded according to the manufacturer’s instructions.

### Statistical analysis

Wilcoxon matched-pairs signed rank test was performed using Graphpad Prism software.

## Results

### Oxazolidinones demonstrated broad activity against Gram-positive endophthalmitis isolates, with tedizolid exhibiting greater potency than linezolid

A representative epsilometer assay is shown in Figure 1A. The overall distribution of minimum inhibitory concentration (MIC) values and descriptive statistics for linezolid and tedizolid is shown in Figure 1B. The median MIC for linezolid was 1.0 µg/ml (range of 0.38-4.0) compared to 0.19 µg/ml (range of 0.064-0.75) for tedizolid. Tedizolid MICs were significantly lower than linezolid MICs by Wilcoxon matched-pairs signed rank analysis (P<0.001), indicating greater potency in vitro.

**Figure 1.**
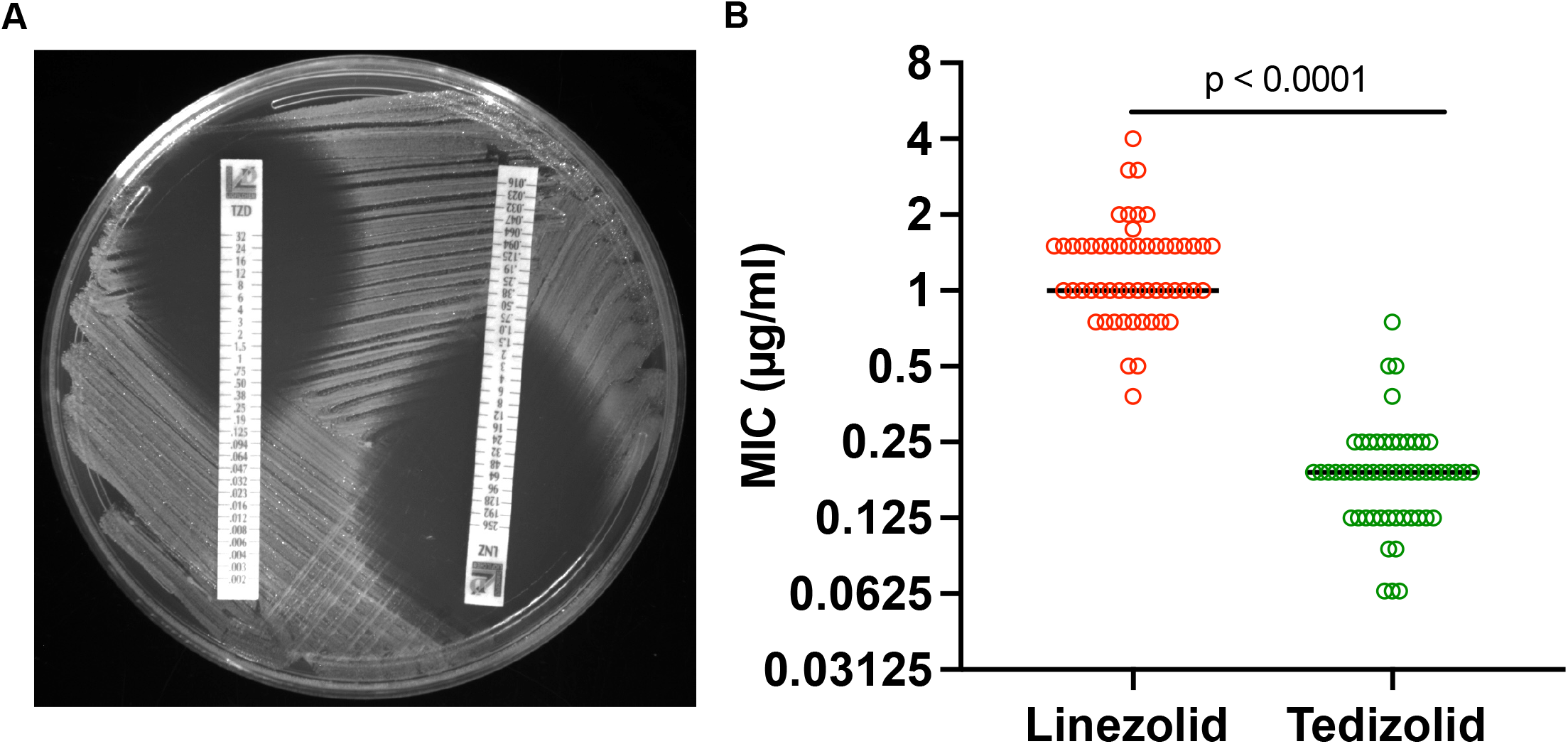
Evaluation of oxazolidinones against endophthalmitis isolates. **A**. Sample plate with a coagulase negative staphylococci isolate tested with epsilometer tests with tedizolid (TZD) and linezolid (LNZ). **B**. MIC values for individual isolates and median are shown. A Wilcoxon matched-pairs signed rank test was used to determine the p-value.

Pathogen-specific results are summarized in Table 1. Using CLSI susceptibility breakpoints, all coagulase-negative staphylococci, *S. aureus, Bacillus* spp,, and *Streptococcus pneumoniae* isolates were susceptible to both agents ^19^. Among *Enterococcus* isolates, one of five was categorized as intermediate to linezolid. No isolate met CLSI criteria for resistance to either oxazolidinone. Using EUCAST criteria, all isolates with available breakpoint interpretations were categorized as susceptible to linezolid ^20^. EUCAST currently does not provide tedizolid breakpoints for *Enterococcus* spp., precluding susceptibility categorization for that organism group.

**Table 1.** Oxazolidinone MICs and susceptibility of Gram-positive endophthalmitis isolates.

| Group<sup>a</sup>, (n-value) | Antibiotic | MIC Median<sup>b</sup> | MIC90<sup>b</sup> | MIC Range<sup>b</sup> | % Susceptible<sup>d</sup> |
| --- | --- | --- | --- | --- | --- |
| CoNS (20) | Linezolid | 1.0 | 2.0 | 0.5-3.0 | 100% |
|  | Tedizolid | 0.19 | 0.25 | 0.064-0.25 | 100% |
| *S. aureus* (18) | Linezolid | 1.5 | 1.5 | 0.75-2.0 | 100% |
|  | Tedizolid | 0.19 | 0.19 | 0.125-0.25 | 100% |
| *Bacillus spp* (10) | Linezolid | 1.0 | 1.0 | 0.38-1.5 | 100% |
|  | Tedizolid | 0.158 | 0.25 | 0.064-0.25 | 100% |
| *Enterococcus spp* (5) | Linezolid | 2.0 | n/a | 1.5-4.0 | 80% |
|  | Tedizolid | 0.5 | n/a | 0.19-0.75 | 80% |
| *S. pneumoniae* (1) | Linezolid | 0.5 | n/a | n/a | 100% |
|  | Tedizolid | 0.064 | n/a | n/a | 100% |

a. CoNS, coagulase negative staphylococci
b. All units µg/ml
c. n/a, not available due to low numbers
d. based on CLSI guidelines

## Discussion

The history of empiric endophthalmitis therapy illustrates that antibiotic selection is influenced by factors beyond antimicrobial spectrum alone. Safety, pharmacokinetics, antimicrobial resistance, and evolving understanding of disease pathogenesis have all shaped current practice. For example, amikacin has largely been replaced by ceftazidime as the Gram-negative covering component of empiric therapy because of concerns regarding toxicity ^21,22^. The present study suggests that oxazolidinones merit similar consideration for treatment of Gram-positive intraocular pathogens.

Here, linezolid and tedizolid demonstrated in vitro activity against Gram-positive endophthalmitis isolates. Tedizolid consistently exhibited lower MIC values than linezolid, with an overall median MIC that was more than five-fold lower. Nearly all isolates were categorized as susceptible using CLSI guidelines, including species associated with rapid progressive and vision-threatening infections such as *Bacillus* spp. and *Enterococcus* spp. The lone non-susceptible isolate demonstrated only a modest elevation in linezolid MIC, suggesting that clinically meaningful resistance to oxazolidinones remains uncommon among Gram-positive endophthalmitis pathogens. Additionally, epsilometer strips are thought to overestimate the MIC values for oxazolidinones compared to the broth-dilution method, so the reported values are likely conservative ^23^.

This study has limitations. The study evaluated in vitro susceptibility but did not assess intraocular pharmacokinetics. This is particularly important for intraocular therapeutics for which the duration of a drug exposure may influence efficacy. For example, moxifloxacin exhibits a short intravitreal half-life, limiting its utility as a primary agent for treatment of bacterial endophthalmitis despite excellent in vitro activity ^24^. Future studies should therefore define ocular pharmacokinetics and duration of exposure above MIC for both linezolid and tedizolid. Retinal toxicity was not examined. Although linezolid has extensive clinical use and excellent systemic bioavailability, its intraocular safety profile following local administration remains undefined.

The findings of this study suggest that oxazolidinones should not necessarily be viewed as replacements for vancomycin, but rather as agents with several potential applications in endophthalmitis management. These may include treatment of infections caused by vancomycin-resistant Gram-positive organisms, incorporation into vancomycin-sparing treatment strategies, and, potentially to attenuate toxin-mediated retinal injury through anti-virulence effects that have been reported for some Gram-positive pathogens. Additional pharmacokinetic, toxicity, and efficacy studies will be required to determine whether these in vitro advantages translate into clinical benefit. Once tested these oxazolidinone antibiotics have the potential to be an important addition to our armamentarium in the treatment against sight threatening endophthalmitis.

## Ethical approval and consent to participate

This work required no institutional ethics review.

## Author contributions

Conceptualization: RMQS, EGR

Data curation: MR, EAY, RMQS

Formal analysis: EAY, EGR, RMQS

Funding acquisition: AWE, RMQS

Investigation: MR, EAY, EGR, RMQS

Writing-original draft: MR, RMQS

Writing-review & editing: MR, EAY, EGR, AWE, RMQS

